# Ascorbate peroxidase neofunctionalization at the origin of APx-R and APx-L: evidences from basal Archaeplastida

**DOI:** 10.1101/2020.08.18.255851

**Authors:** Fernanda Lazzarotto, Paloma Koprovski Menguer, Luiz-Eduardo Del-Bem, Márcia Margis-Pinheiro

**Affiliations:** Programa de Pós-graduação em Biologia Celular e Molecular, Centro de Biotecnologia, Universidade Federal do Rio Grande do Sul, Porto Alegre, Brazil; Departamento de Botânica, Instituto de Ciências Biológicas, Universidade Federal De Minas Gerais, Belo Horizonte, Brazil; Programa de Pós-Graduação em Genética e Biologia Molecular, Departamento de Genética, Universidade Federal do Rio Grande do Sul, Porto Alegre, Brazil

**Keywords:** APx, APx-R, APx-L, catalytic sites, substrate, divergence

## Abstract

Ascorbate peroxidases (APx) are class I members of the non-animal peroxidases superfamily, a large group of evolutionarily related enzymes. Through mining in public databases, our group has previously identified two unusual subsets of APx homologs, disclosing the existence of two uncharacterized families of class I peroxidases, which were named ascorbate peroxidase-related (APx-R) and ascorbate peroxidase-like (APx-L). As APx, APx-R proteins possess all catalytic residues required for peroxidase activity. Nevertheless, these proteins do not contain residues known to be critical for ascorbate binding, implying that members of this family must use other substrates while reducing hydrogen peroxide. On the other hand, APx-L proteins not only lack ascorbate-binding residues, as do not contain any residue known to be essential for peroxidase activity, in contrast with every other member of the non-animal peroxidase superfamily, which is composed by over 10,000 proteins distributed among bacteria, archaea, fungi, algae, and plants. Through a molecular phylogenetic analysis performed with sequences derived from basal Archaeplastida, we now show the existence of hybrid proteins, which combine features of these three families. Analysis performed on public databases show that the prevalence of these proteins varies among distinct groups of organisms, accounting for up to 33% of total APx homologs in species of green algae. The analysis of this heterogeneous group of proteins sheds light on the origin of APx-R and APx-L, through a process characterized by the progressive deterioration of ascorbate-binding sites and catalytic sites towards neofunctionalization.

## 1. Introduction

Ascorbate peroxidases (APx) (EC 1.11.1.11) are enzymes that catalyze the hydrogen peroxide (H_2_O_2_) reduction in oxygenic photosynthetic organisms in a process dependent on ascorbate. As for other heme-containing enzymes, the catalytic mechanism involves the formation of an oxidized Compound I intermediate and its subsequent reduction by the substrate, in this particular case ascorbate, in two sequential electron transfer steps [1,2]. The catalytic mechanism, crystal structure, and ascorbate binding of APx have been extensively studied in the past thirty years [3–9]. Through crystallographic information and site-directed mutagenesis, it was possible to dissect the main residues implicated in APx activity, which are allocated in two structural domains surrounding the heme[10]. The C-terminal domain contains the proximal histidine (His163), which is hydrogen-bonded to the heme moiety. The N-terminal domain harbors the distal histidine (His42) and an arginine (Arg38) that are crucial for the heterolytic cleavage of H_2_O_2_, as well as a tryptophan residue (Trp41) implicated with heme-binding [10–13]. The interaction of APx with ascorbate is only possible through the existence of an arginine located nearby the proximal histidine (Arg172), since mutations in this residue completely abolished APx activity towards this substrate [6][8]. Nevertheless, N-terminal lysine (Lys30) and cysteine (Cys32) residues also seem to contribute to ascorbate binding, but to a much lesser extent [7].

The APx family is part of the non-animal peroxidase superfamily, a group of evolutionarily related enzymes distributed among bacteria, algae, fungi, and plants [14]. Despite the low sequence homology among superfamily members, proteins that belong to this group share features like folding, secondary structure, and catalytic domains. Molecular phylogeny analyses separate the superfamily members in three well-supported classes, among which over ten families are accommodated [10,15–17]. Through mining in public databases, our group has identified an unusual subset of APx proteins that did not cluster with classic APx members in phylogenetic analyses [18]. These proteins show several conserved substitutions when compared to classic APx, which include the substitution of Trp41 by a phenylalanine, in resemblance of other superfamily peroxidases, and the lack of Arg172, suggesting that they should not be able to use ascorbate as a catalytic substrate. After a refined phylogenetic analysis, we were able to confirm that these proteins constitute a distinct family of heme peroxidases, which was called ascorbate peroxidase-related (APx-R). Members of this new family are mostly predicted to contain a chloroplast transit peptide and to be encoded by a single-copy gene in virtually all plant genomes [18]. Functional analysis demonstrated that *APx-R* knockdown disturbed the redox metabolism in rice (*Oryza sativa* L.) while analyses conducted in arabidopsis knockout plants revealed the importance of this peroxidase to oxidative protection and hormonal steadiness in seeds [18,19]. More recently, through a robust phylogenetic analysis, we were able to classify APx-R as a class I member of the non-animal peroxidases superfamily, along with APx, catalase-peroxidase (KatG), cytochrome-c peroxidase (CcP) and APx-CcP families [17]. Moreover, this analysis also revealed the existence of another well-supported family composed by a small subset of proteins, currently annotated as APx homologs, which was named ascorbate peroxidase-like (APx-L). In contrast to what was observed for APx-R, APx-L proteins do not harbor any of the catalytic residues described as key for hydrogen peroxide enzymatic removal. Arabidopsis APx-L, also named as TL-29 (for thylakoid lumen 29kDa protein) or APx04, has been functionally and structurally investigated previously [20–22]. Purification of native APx-L from chloroplasts extract have showed that this protein accounts for one of the most abundant in the thylakoid lumen of arabidopsis [21], while recombinant expression has confirmed that APx-L is not able to bind the heme, ascorbate or catalyze hydrogen peroxide removal [20]. Interestingly, analyses conducted in arabidopsis knockout plants suggest that APx-L is functional and participates on photosystem protection and seed coat formation, through mechanisms which are yet unclear [22]. Despite the strong support for their reclassification as new families, APx-R and APx-L have been continuously addressed as ascorbate peroxidases in the literature, causing the misinterpretation of experimental data and impairing the discussion over H_2_O_2_ scavenging metabolism in plants. To understand how these three families are distributed among basal organisms in the plant lineage and to access the origin of the structural and functional diversity among them, we performed a comprehensive phylogenetic analysis using sequences retrieved from species of red and green algae and bryophytes. As expected, tree topology showed three well-supported clusters indicating a clear separation between APx, APx-R and APx-L families. However, sequences that compose each cluster proved to be more heterogeneous than expected, and hybrid proteins, which combine features from more than one family, are now described for the first time. The acknowledgment of this diversity provides the basis to explain the origin of APx-R and APx-L from ascorbate peroxidases. From this perspective, we speculate that the acquisition of new functions must have coordinated with the loss of APx activity in these proteins, culminating with their establishment as distinct families in photosynthetic organisms.

## 2. Results

### 2. 1. Hybrid proteins share characteristics of distinct families

Phylogenetic analysis was carried out with 309 protein sequences from species of algae and non-vascular plants, in addition to 39 sequences encoding KatG and CcP, which were used as outgroups. The dataset included 118 sequences previously deposited in RedOxiBase annotated either as APx (which encompass both APx and misannotated APx-L) or APx-R, and 191 sequences from charophytes derived from 39 assembled transcriptomes (http://www.onekp.com/public_data.html) and the complete genome of *Klebsormidium nitens* (http://www.plantmorphogenesis.bio.titech.ac.jp/~algae_genome_project/klebsormidium/). From this analysis, it is possible to distinguish five main clusters, three of which corresponding to APx (purple), APx-R (orange) and APx-L(cyano) families, in addition to KatG and CcP (Fig. 1). The tree topology is in agreement with our previous study, showing that APx family is more closely related to KatG and CcP than to APx-R and APx-L (Fig. 1), although being all part of the same peroxidase class [17]. A detailed analysis of each group revealed the existence of sequence variants, which are marked in red (Fig. 1). These sequences consist of proteins that present characteristics that are singular of at least two different families (e.g., the simultaneous occurrence of Phe41, an APx-R feature, and Arg172, an APx specific residue), what has not been observed previously [17]. From now on we will name these proteins as hybrids.

**Figure 1.**
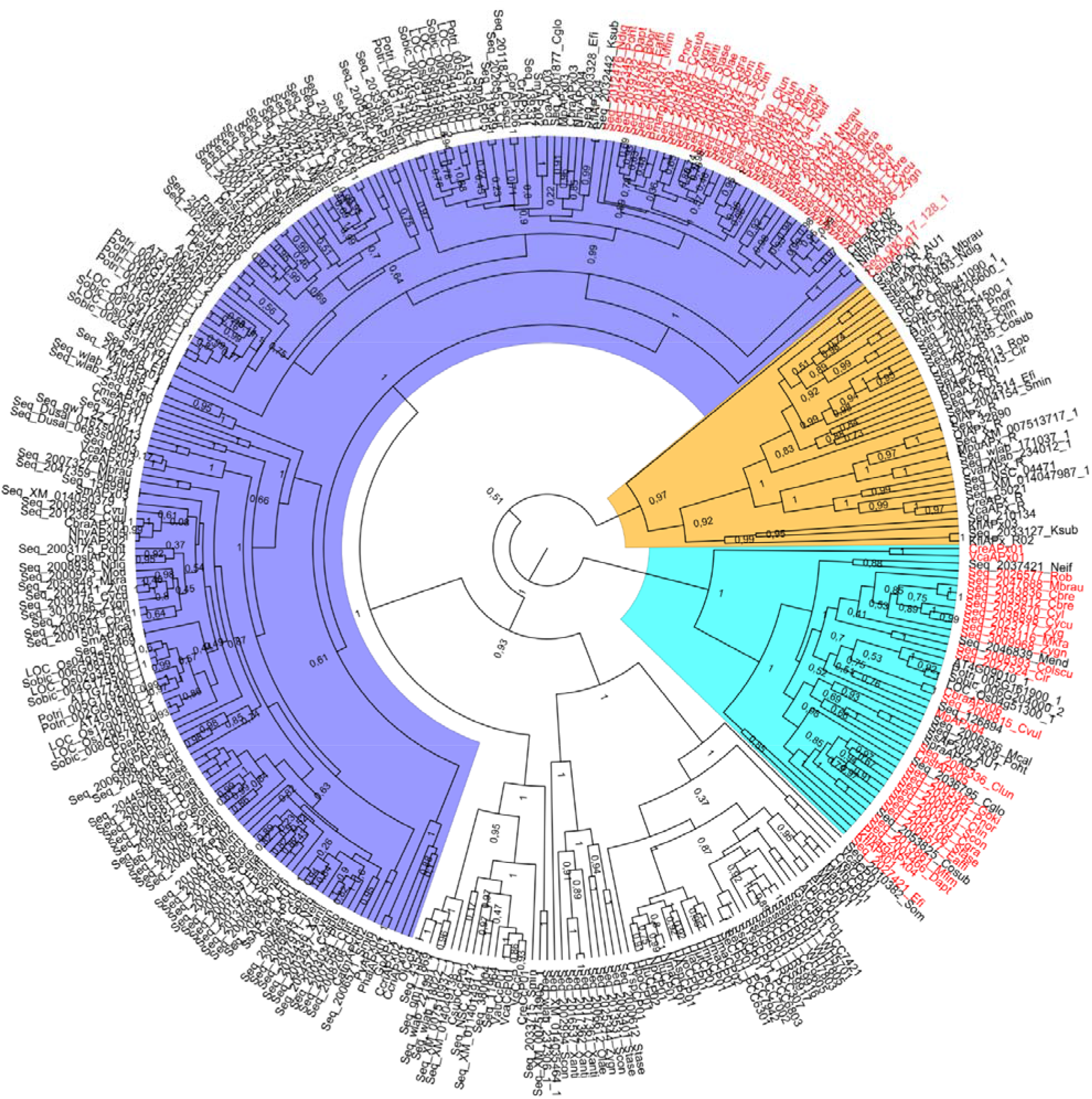
The phylogenetic relationship between APx, APx-R, APx-L, CcP and CP was reconstructed using the Bayesian method. A total of 348 protein sequences were included in the analysis, and ambiguous positions were removed from the alignment. APx, APx-R and APx-L families separated in three well-supported clusters and these are colored in purple (APx), orange (APx-R) and cyano (APx-L). Protein sequences that display hybrid features are indicated in red. The posterior probabilities are discriminated above each branch.

### 2. 2. Hybrid proteins are more prevalent in species of green algae

To analyze the prevalence of hybrid proteins in different groups of organisms, we decided to look for variants in APx/APx-L/APx-R sequences in public databases. For this purpose, 493 protein sequences were retrieved from RedOxiBase and further classified into rhodophytes, chlorophytes, charophytes, bryophytes, gymnosperms, and angiosperms. In addition, 143 charophytes sequences from our dataset were also evaluated, adding up to 636 sequences. The presence or absence of key residues implicated in catalytic activity and ascorbate binding determined their categorization, which followed the protein signature described for each family [23]. To be considered an APx, proteins should present the catalytic residues Arg38, Trp41, His42 and His163; in addition to the ascorbate-binding residue Arg172. For APx-R classification, proteins should include Arg38, Phe41, His42 and His163 and position 172 should be occupied by any residue other than arginine. To be classified as APx-L, all the above-mentioned positions should be occupied by other residues (Fig. 2). Proteins were considered hybrids when exhibited different arrangements of amino acids in these positions. The occurrence of each group of proteins in the analyzed classes is summarized in figure 3, and their prevalence by species is provided in table S1. From this analysis, it is possible to infer that the diversification of APx must have occurred at the basis of Viridiplantae. However, one must consider that the low number of sequences currently available from rhodophytes could be leading to a biased interpretation in this subject. In Viridiplantae, hybrid proteins are considerably more prevalent in organisms belonging to deep-branching clades, accounting for up to 33% of total analyzed sequences derived from charophytes. While APx-R is present in all classes, APx-L could only be detected in charophytes, bryophytes, and angiosperms, in rates from 3% to 9%. Among the analyzed groups, gymnosperms are the organisms with the higher prevalence of APx proteins (93%), followed by angiosperms. All complete protein sequences deposited in RedOxiBase and classified as APx-R, APx-L or hybrids are listed in table S2. Although hybrid proteins are more prevalent in basal organisms, what is partly explained by the recent expansion of the APx family (table S1), a few can be found in vascular plants. An interesting example is observed in poplar (*Populus trichocarpa*), in which a single gene encodes an APx and a hybrid protein through alternative splicing, producing a variant protein that potentially retained the ability to bind ascorbate while deprived of peroxidatic activity (Fig. 4).

**Figure 2.**
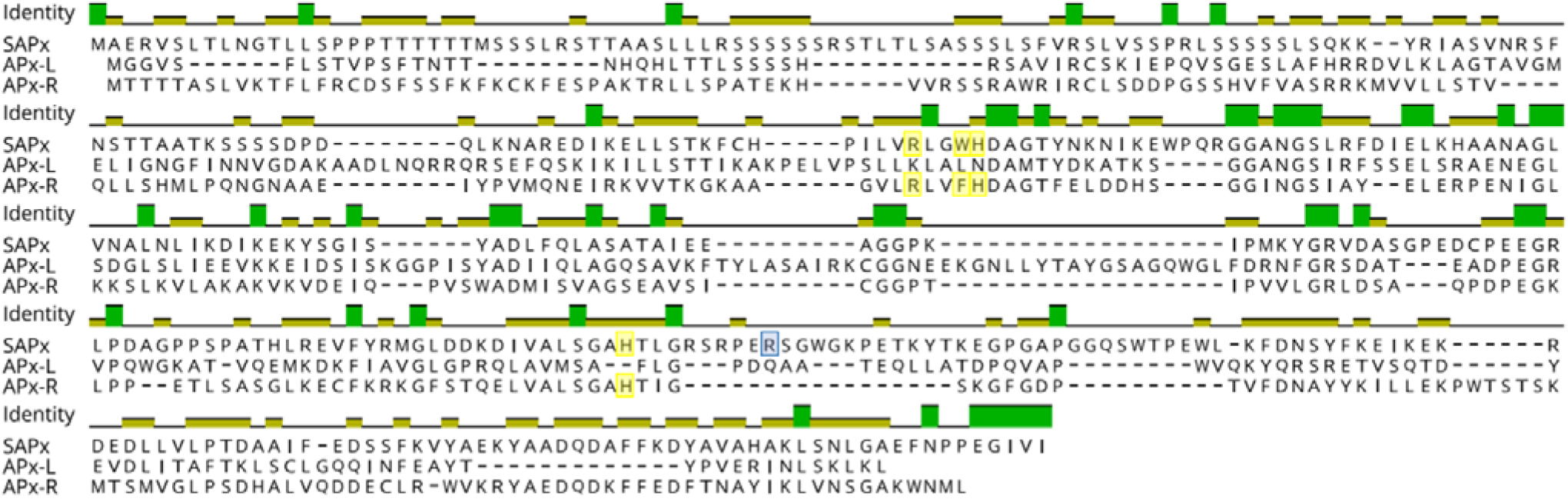
Protein alignment of Arabidopsis stromal APx (SAPx; At4g08390), APx-R (At4g32320) and APx-L (At4g09010). Positions related to APx catalytic activity are highlighted in yellow and to ascorbate binding, in blue.

**Figure 3.**
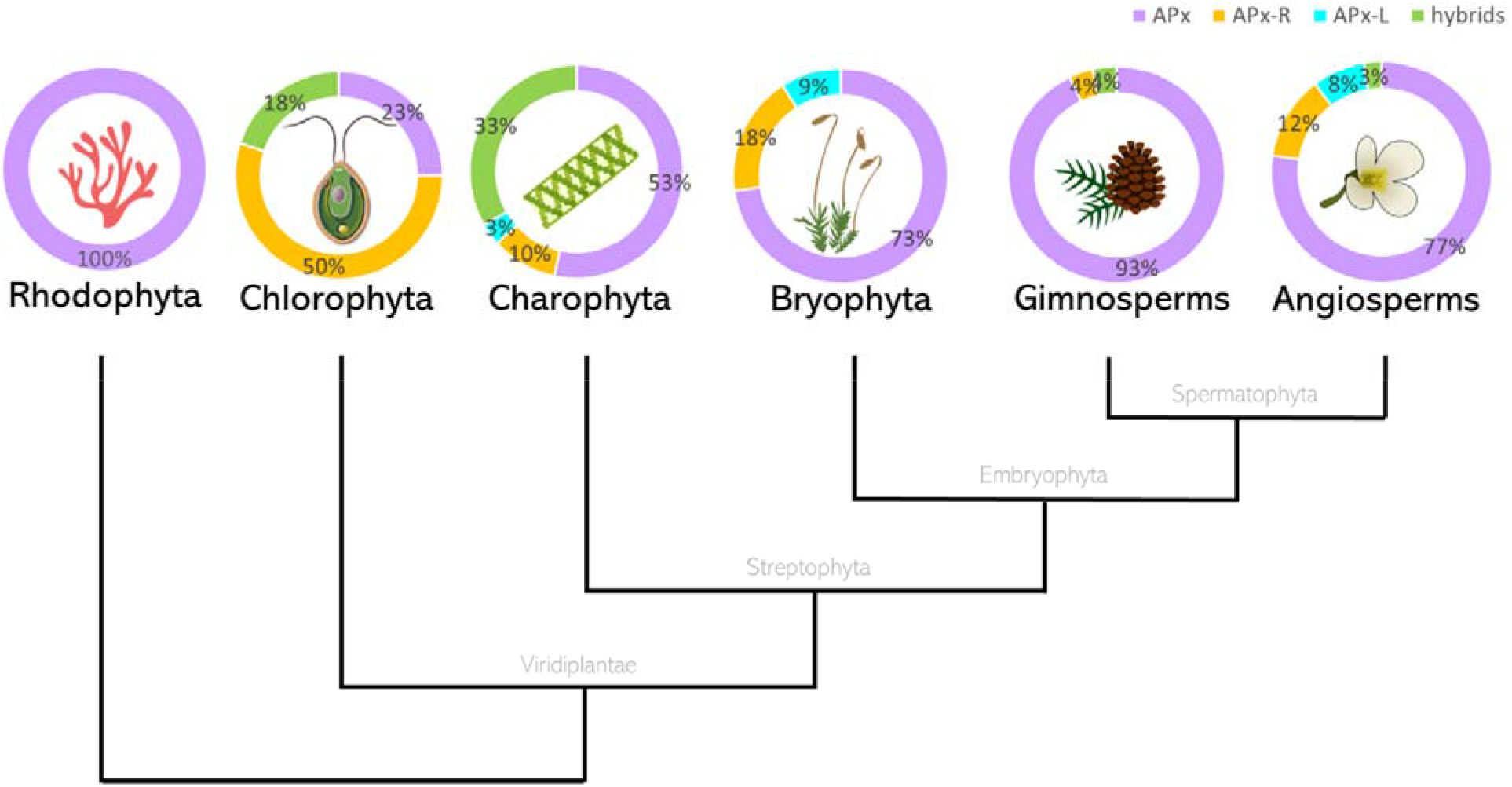
Protein sequences currently annotated in RedOxiBase were classified according to key residues as APx, APx-R, APx-L or hybrids. The total number of sequences analyzed is 6, 22, 177, 11, 28, and 389, respectively. Each group of proteins is represented by the following colours: APx, purple; APx-R, orange; APx-L, cyano; and hybrids are shown in green. *The data presented for Charophytes include sequences currently annotated in RedOxiBase in addition to 147 assembled sequences obtained from transcriptomic data, available in table S3.

**Figure 4.**
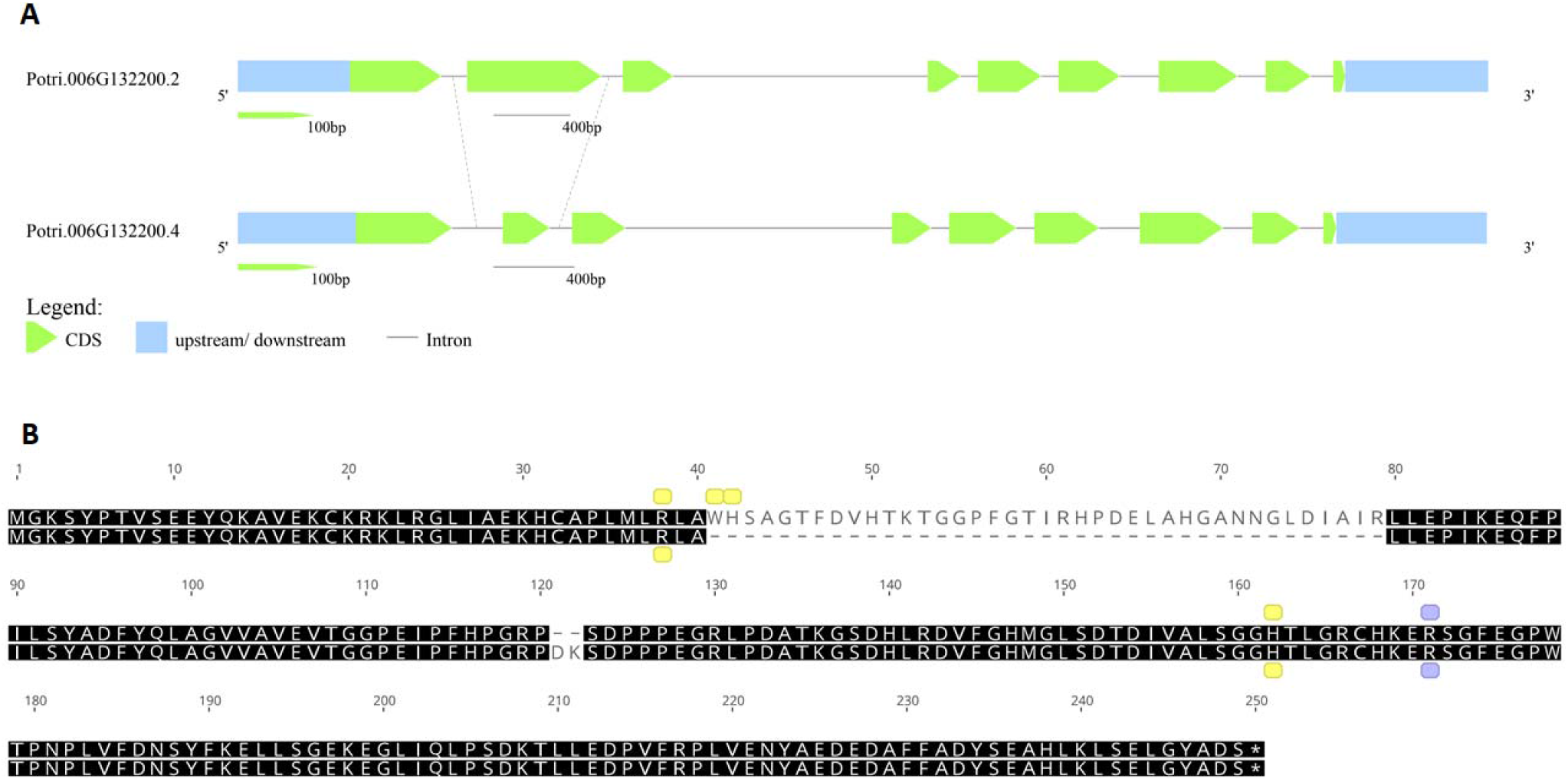
Alternative splicing favours the production of an APx or a hybrid protein in *Populus trichocarpa*. The gene Potri.006G132200 encodes four transcripts, two of which differ on the size of the second exon (A). The retention of the truncated version of the exon causes the loss of two catalytic residues, which are essential for peroxidase activity (Trp41 and His42). Ascorbate binding arginine is indicated in blue and catalytic residues in yellow (B).

### 3. Point mutations and small deletions may have resulted in the emergence of APx-R, APx-L and hybrids

To assess the extent of variation found in these distinct groups of proteins, we performed an alignment using sequences derived from chlorophytes and charophytes, which is presented in figure 5. Through this analysis, it is possible to observe that point mutations and two small deletions around the proximal histidine are the main cause of variation found in these families regarding catalytic and ascorbate-binding residues. This analysis also suggests that hybrid proteins accumulated fewer mutations than APx-R and APx-L, resembling intermediates among families.

**Figure 5.**
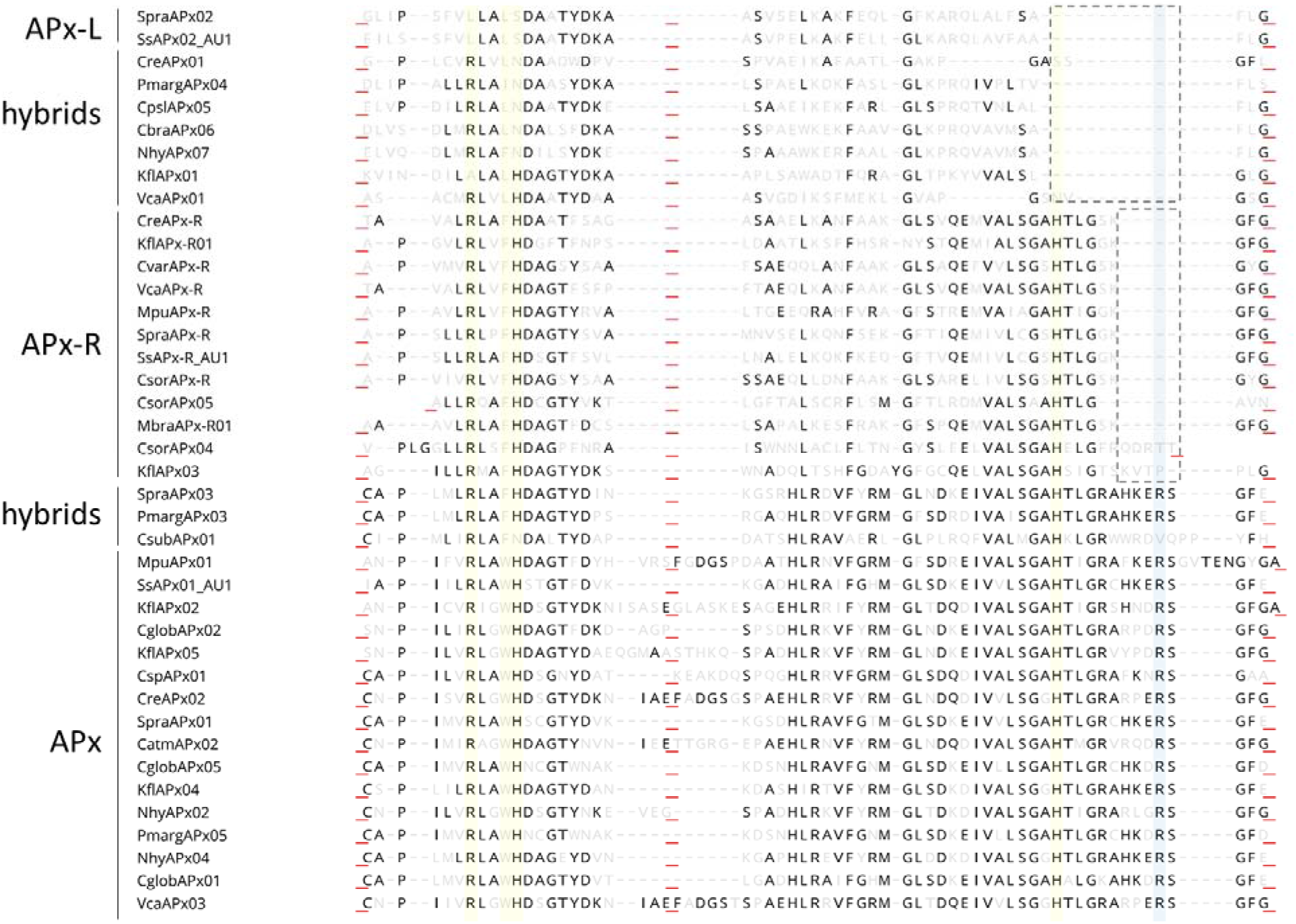
Point mutations and two small deletions are involved with APx-R and APx-L differentiation. Sequences of each protein category were retrieved from RedOxiBase and domains that contain relevant residues are shown. Conserved residues are shown in black and others, in light grey. Positions implicated with catalytic activity are showed in yellow and with ascorbate-binding, in blue. While point mutations may be responsible for the progressive loss of N-terminal catalytic residues, two small deletions appear to be the main cause leading to Arg172 and His163 loss. Red underscores evidence domains collapsing for better visualization.

A large proportion of the hybrid proteins identified in this study is organized in a cluster that integrates the well-supported APx group in our phylogenetic analysis (Fig. 1). Despite the overall similarity with APx, these charophyte sequences contain a phenylalanine at position 41 instead of the APx-conserved tryptophan, which suggests that this mutation could have been implicated with the first modifications that accompanied the establishment of APx-R (Fig. 6). Similarly, two charophyte sequences that also integrate the APx group exhibit an arrangement of residues that might help explaining the evolution of APx-L. These sequences also contain phenylalanine at position 41, while have lost the distal histidine and the ascorbate-binding arginine. These two types of sequences are here named as proto-APx-R and proto-APx-L, respectively, and their identification provides support for the establishment of APx-R and APx-L families from APx diversification.

**Figure 6.**
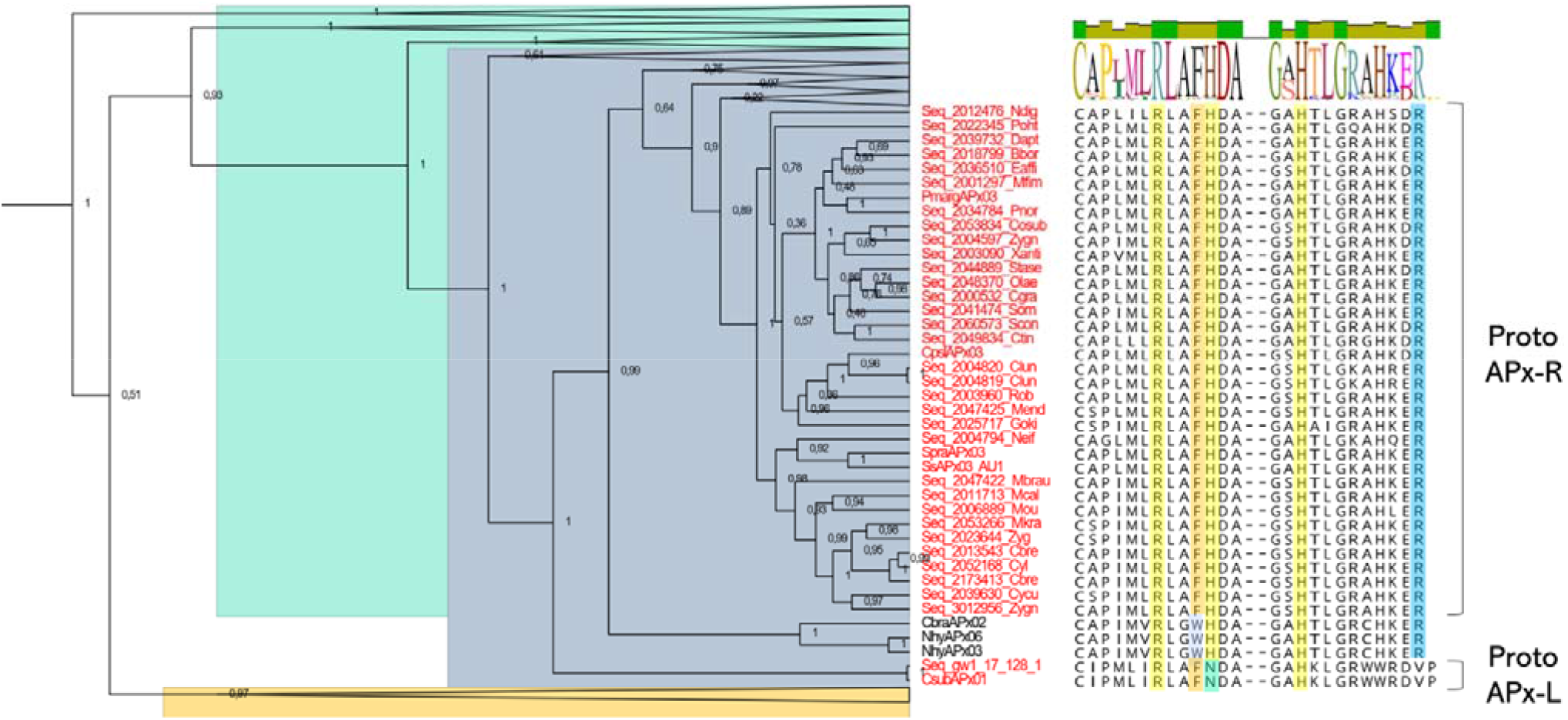
Detailed analysis of a clade from APx cluster. Hybrid sequences derived from Charophytes exhibit a single mutation from tryptophan to phenylalanine at position 41, being therefore addressed as proto-APx-R. The loss of arginine at position 172 and the replacement of the distal histidine by an asparagine illustrate the diversification pathway towards APx-L establishment, by which reason these sequences are referred to as proto-APx-L. APx group is showed in blue; APx-R, in orange; and APx-L, in green. All other clades showed in figure 1 were collapsed for better visualization.

## 3. Discussion

The structure and function of ascorbate peroxidase have been extensively studied since recombinant expression and site-directed mutagenesis techniques were established. Due to its overall homology to CcP, ascorbate peroxidase was expected to display a similar enzymatic activity. However, because APx catalysis, as well it is for other heme peroxidases, relies on the formation of a porphyrin-based radical and not on a protein-based radical, which is the case for CcP, APx quickly became an interesting model to dissect heme peroxidase catalysis in non-animal organisms [24].

Since the publication of APx first crystal [3] and later of APx-ascorbate complex structure [6], all main residues implicated in the enzymatic activity displayed towards this substrate were identified. Site-directed mutagenesis have later confirmed that two histidines (His42 and His163), an arginine (Arg38) and a tryptophan (Trp41) are essential for APx to catalyze hydrogen peroxide scavenging [9,25–27]. Apart from Trp41, which was successfully substituted by phenylalanine in proteins belonging to classes II and III [15,28], all the other residues are conserved and have proven to be critical for peroxidases to interact with the heme moiety and with hydrogen peroxide.

Regarding APx interaction with its substrate, despite the evidence that Lys30 and Cys32 could be involved, an arginine at position 172 has proven to be the critical residue for APx to bind ascorbate. While substitutions at this position have abolished APx activity towards this substrate, they did not interfere with APx binding to non-physiological aromatic substrates *in vitro*, indicating that APx might could also interact with other molecules *in vivo* through distinct sites [7,29]. Because Arg172 is missing in APx-R, we have previously suggested that members of this family might display enzymatic activity using other substrates that not ascorbate, which was recently confirmed through recombinant expression and enzymatic analysis from purified protein (unpublished). Nevertheless, the natural substrate for APx-R *in vivo* remains to be experimentally determined. Interestingly, this is not the case for APx-L. It has been previously reported that arabidopsis APx-L is devoid of catalytic activity and unable to bind ascorbate or the heme, although being expressed at high levels and somehow functional, with knockout plants showing a chlorotic phenotype and producing seeds with reduced longevity [17,20–22]. In addition, authors have showed experimental evidence of APx-L association with the photosystem II, supporting a divergent role for this protein. In this scenario, we hypothesize that the acquisition of new functions must have been accompanied by the progressive loss of the ancestral and eventually obsolete ascorbate-binding site in APx-R, and catalytic sites, in APx-L.

It is believed that the non-animal peroxidase superfamily, which is composed of twelve families of peroxidases accommodated in three classes, is derived from an ancestral peroxidase found in an ancient prokaryote [15,17]. The superfamily emergence is explained as the result of countless duplication events, and it seems to be tightly associated with mitochondria and chloroplast acquisitions. Because of this, all superfamily enzymes share characteristics like structure and folding, despite differences observed in amino acid composition [10]. It is also likely that the accumulation of point mutations eventually culminated in the establishment of the families that we encounter nowadays. In this scenario, the identification of hybrid proteins is outstanding evidence of this evolutionary process.

The data generated in this study culminated in the proposition of a model to explain the emergence of these proteins in autotrophic eukaryotes, which is presented in figure 7. For being more closely related to bacterial KatG and other class I enzymes, APx is believed to be more ancestral than APx-R and APx-L. Therefore, the identification of hybrid proteins suggests that the establishment of these two families may have been accompanied by the progressive deterioration of APx catalytic and/or ascorbate binding sites. This hypothesis is also sustained on the identification of proteins that maintain ascorbate peroxidases features while having the APx-R/classII/classIII-specific phenylalanine at position 41 (proto-APx-R), or of proteins lacking distal histidine while keeping others catalytic residues (proto-APx-L). The absence of typical APx-L in chlorophytes indicate that this group of proteins established more recently in a Streptophyta ancestor; however, this hypothesis remains to be confirmed when genomic and transcriptomic data from other species that compose this group becomes available.

**Figure 7.**
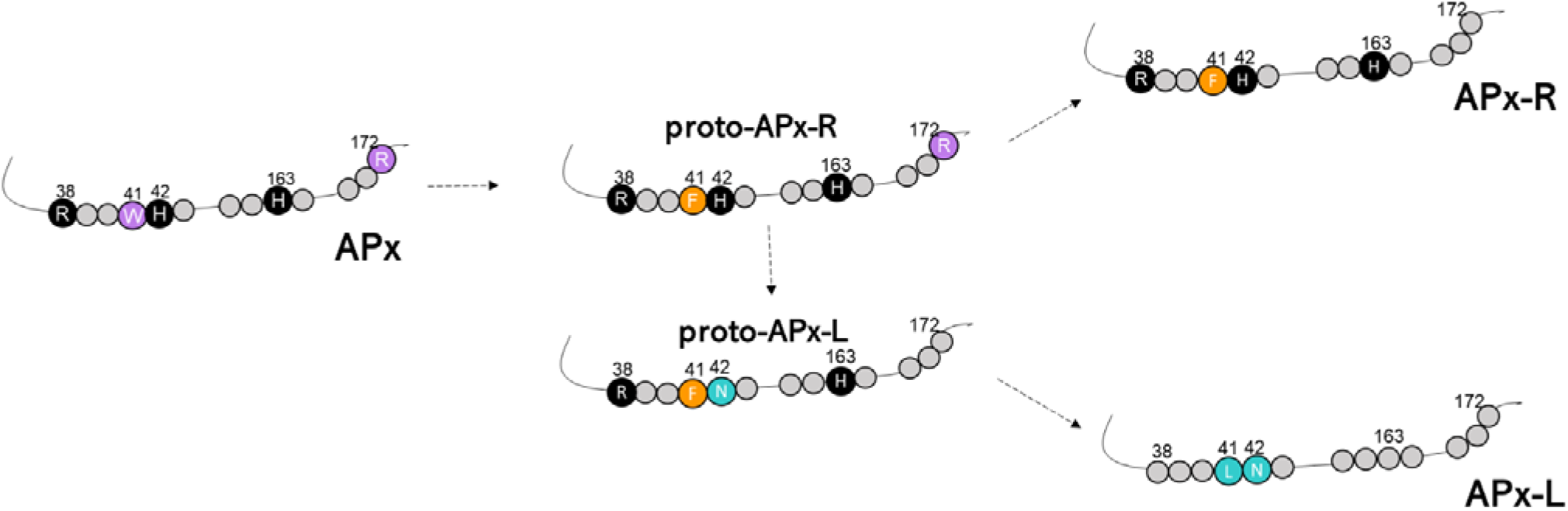
The proposed model to explain the origin of APx-R and APx-L. Relevant residues are represented, and their position is discriminated. Conserved amino acids are indicated and coloured according to the group they are primarily found.

The functionality of APx-L and hybrid proteins is yet unclear. While in the first case, there is a homogeneity of characteristics that is observed for most family members (e.g., amino acid composition, exon-intron structure, presence of a transit peptide), hybrid proteins present a myriad of amino acid arrangements, which might be leading to somewhat dissimilar functions. While some proteins may have retained the ability to bind ascorbate while lacking peroxidatic activity, others may function in another direction. The characterization of APx-R and APx-L function *in vivo* will provide the basis to understand which are the key residues implicated with their role in the antioxidant metabolism, and how hybrids might behave in this context. The dissimilar distribution of each family and hybrid proteins in the analyzed groups also suggests that this process occurred under distinct selective pressures in aquatic and terrestrial organisms. Nevertheless, the maintenance of hybrid proteins must have been advantageous to some plants, as in the case of poplar. Despite the limited number of sequences from basal organisms, an interesting observation is that a few algae species contain more than one APx-R encoding gene, in contrast to what we have observed previously for vascular plants, in which genomes the duplication of this gene seems to be detrimental [18]. In conclusion, this work provides new evidence on the origin and evolution of APx, APx-R and APx-L families and points out to the existence of significant diversity in the antioxidant metabolism, which should be taken into consideration when accessing ascorbate peroxidase and antioxidant metabolism function in photosynthetic organisms.

## 4. Methods

### 4. 1. Sequence retrieval

Publicly available protein sequences used in this study were retrieved from RedOxiBase [30]. For charophytes, the sequences were retrieved from the OneKP data (http://www.onekp.com/public_data.html) and the complete predicted proteome of *Klebsormidium nitens* (http://www.plantmorphogenesis.bio.titech.ac.jp/~algae_genome_project/klebsormidium/) using blastp searches (e-value < e^−5^) with the known *Arabidopsis thaliana* proteins as queries. Redundant sequences were eliminated using in-house scripts, keeping only the longest protein sequence. Sequences that were shorter than 50% of the *Arabidopsis* query, were regarded as incomplete and discarded.

### 4. 2. Sequence alignment and phylogenetic analysis

Sequence alignments were conducted using MUSCLE algorithm [31] with default parameters available at MEGA 7.0 (Molecular Evolutionary Genetics Analysis) [32]. The phylogenetic tree was reconstructed using protein sequences of conserved domains by Bayesian inference using BEAST2.4.5 [33]. The best fit model of amino acid replacement was LG with invariable sites and gamma-distributed rates, which was selected with ProtTest [34]. The Birth and Death Model was selected as tree prior, and 50,000,000 generations were performed with Markov Chain Monte Carlo algorithm (MCMC) [35] for the evaluation of posterior distributions. After manual inspection of the alignment, 348 sequences and 233 sites were used in the analysis. Convergence was verified with Tracer [36], and the consensus tree was generated using TreeAnnotator, available at BEAST package. The resulting tree was analyzed and edited using FigTree v.1.4.3 (http://tree.bio.ed.ac.uk/software/figtree). Figures that show sequence alignments were generated using Geneious Prime 2020.1.1 (https://www.geneious.com); for the gene structure figure, GSDS 2.0 was employed [37].

## Abbreviations

APx: Ascorbate peroxidase
APx-R: Ascorbate peroxidase-related
APx-L: Ascorbate peroxidase-like
KatG: Catalase peroxidase
CcP: Cytochrome-c peroxidase
H_2_O_2_: Hydrogen peroxide

## Supplementary material

**Table S1.** Occurrence of APx, APx-R, APx-L and hybrid proteins in species of Archeaplastida based on sequences currently deposited in RedOxiBase and OneKP.

|  | APx | APx-R | APx-L | Hybrid |
| --- | --- | --- | --- | --- |
| **Rhodophyta** |  |  |  |  |
| *Cyanidioschyzon merolae* | 1 | 0 | 0 | 0 |
| *Galdieria partita* | 1 | 0 | 0 | 0 |
| *Galdieria sulphuraria* | 2 | 0 | 0 | 0 |
| *Porphyra raitanensis* | 1 | 0 | 0 | 0 |
| *Porphyra yezoensis* | 1 | 0 | 0 | 0 |
| **Chlorophyta** |  |  |  |  |
| *Chlamydomonas reinhardtii* | 1 | 1 | 0 | 1 |
| *Chlorella sorokiniana* | 1 | 5 | 0 | 0 |
| *Chlamydomonas sp* | 1 | 0 | 0 | 0 |
| *Coccomyxa subellipsoidea* | 0 | 0 | 0 | 1 |
| *Chlorella variabilis* (C.sp. NC64A) | 0 | 1 | 0 | 0 |
| *Micromonas pusilla* | 1 | 1 | 0 | 0 |
| *Ostreococcus lucimarinus* | 0 | 1 | 0 | 0 |
| *Ostreococcus tauri* | 1 | 1 | 0 | 0 |
| *Volvox carteri* | 1 | 1 | 0 | 1 |
| **Charophyta** |  |  |  |  |
| *Chlorokybus atmophyticus* | 3 | 1 | 0 | 0 |
| *Chara braunii* | 2 | 0 | 0 | 1 |
| *Chaetosphaeridium globosum* | 3 | 0 | 0 | 0 |
| *Closterium peracerosum-strigosum-littorale* | 0 | 0 | 0 | 1 |
| *Klebsormidium flaccidum* | 3 | 3 | 0 | 1 |
| *Mesotaenium braunii* | 0 | 1 | 0 | 0 |
| *Nitella hyalina* | 5 | 0 | 0 | 1 |
| *Penium margaritaceum* | 1 | 0 | 0 | 2 |
| *Spirogyra pratensis* | 1 | 1 | 1 | 1 |
| *Spirogyra sp* | 2 | 1 | 1 | 1 |
| *Entransia fimbriat* | 3 | 1 | 0 | 1 |
| *Cosmarium tinctum* | 3 | 1 | 0 | 2 |
| *Desmidium aptogonum* | 2 | 0 | 0 | 2 |
| *Closterium lunula* | 0 | 0 | 0 | 3 |
| *Chaetosphaeridium globosum* | 2 | 0 | 1 | 0 |
| *Netrium digitus* | 3 | 1 | 0 | 1 |
| *Klebsormidium subtile* | 1 | 1 | 0 | 0 |
| *Xanthidium antilopaeum* | 2 | 0 | 0 | 1 |
| *Onychonema laeve* | 1 | 0 | 0 | 1 |
| *Euastrum affine* | 1 | 0 | 0 | 2 |
| *Mesotaenium caldariorum* | 2 | 0 | 1 | 1 |
| *Staurostrum sebaldi* | 2 | 0 | 0 | 2 |
| *Cylindrocystis cushleackae* | 2 | 0 | 0 | 2 |
| *Gonatozygon kinahanii* | 2 | 0 | 0 | 2 |
| *Nucleotaenium eifelense* | 3 | 0 | 1 | 1 |
| *Micrasterias fimbriata* | 1 | 0 | 0 | 2 |
| *Zygnemopsis sp.* | 2 | 0 | 0 | 2 |
| *Cosmarium granatum* | 2 | 0 | 0 | 2 |
| *Pleurotaenium trabecul* | 1 | 0 | 0 | 0 |
| *Chara vulgaris* | 2 | 0 | 0 | 1 |
| *Mesotaenium kramstei* | 2 | 0 | 0 | 2 |
| *Spirotaenia minuta* | 2 | 1 | 0 | 0 |
| *Coleochaete irregularis* | 2 | 1 | 0 | 1 |
| *Bambusina borneri* | 1 | 0 | 0 | 1 |
| *Phymatodocis nordstedtiana* | 2 | 1 | 0 | 2 |
| *Staurodesmus omearii* | 2 | 1 | 0 | 1 |
| *Planotaenium ohtanii* | 2 | 0 | 1 | 1 |
| *Zygnema sp.-A* | 2 | 0 | 0 | 1 |
| *Spirotaenia sp.* | 1 | 0 | 0 | 0 |
| *Cylindrocystis sp.* | 2 | 0 | 0 | 2 |
| *Coleochaete scutata* | 2 | 0 | 0 | 1 |
| *Staurodesmus convergens* | 2 | 0 | 0 | 2 |
| *Mesotaenium endlicherianum* | 0 | 0 | 0 | 2 |
| *Cosmarium subtumidum* | 3 | 1 | 0 | 1 |
| *Zygnema sp.-B* | 2 | 0 | 0 | 2 |
| *Mesotaenium braunii* | 2 | 1 | 0 | 2 |
| *Roya obtusa* | 2 | 1 | 0 | 2 |
| *Cylindrocystis brebissonii* | 2 | 0 | 0 | 4 |
| *Mougeotia sp.* | 2 | 0 | 0 | 1 |
| **Bryophyta** |  |  |  |  |
| *Marchantia paleacea* | 1 | 0 | 0 | 0 |
| *Nothoceros aenigmaticus* | 1 | 0 | 0 | 0 |
| *Physcomitrella patens* | 3 | 1 | 0 | 0 |
| *Marchantia polymorpha* | 3 | 1 | 0 | 1 |
| **Gymnosperms** |  |  |  |  |
| *Pinus pinaster* | 2 | 0 | 0 | 0 |
| *Cycas rumphii* | 2 | 0 | 0 | 0 |
| *Welwitschia mirabilis* | 2 | 0 | 0 | 0 |
| *Zamia fischeri* | 2 | 0 | 0 | 0 |
| *Picea glauca* | 2 | 0 | 0 | 0 |
| *Picea sitchensis* | 2 | 0 | 0 | 0 |
| *Cryptomeria japonica* | 2 | 0 | 0 | 0 |
| *Pinus taeda* | 5 | 0 | 0 | 0 |
| *Ginkgo biloba* | 4 | 1 | 0 | 1 |
| *Picea abies* | 3 | 0 | 0 | 0 |
| *Wollemia nobilis* | 1 | 0 | 0 | 0 |
| **Monocots** |  |  |  |  |
| *Acorus americanus* | 1 | 0 | 0 | 0 |
| *Allium cepa* | 3 | 0 | 0 | 0 |
| *Asparagus officinalis* | 1 | 0 | 0 | 0 |
| *Brachypodium distachyon* | 7 | 1 | 1 | 0 |
| *Crocus sativus* | 1 | 0 | 0 | 0 |
| *Elaeis guineensis* | 2 | 0 | 0 | 0 |
| *Festuca arundinacea* | 0 | 1 | 0 | 0 |
| *Hordeum vulgare* | 7 | 1 | 0 | 0 |
| *Oryza brachyantha* | 0 | 1 | 0 | 0 |
| *Oryza sativa ssp japonica cv Nipponbare* | 9 | 1 | 1 | 0 |
| *Oryza sativa (indica cultivar-group)* | 0 | 1 | 0 | 0 |
| *Pennisetum americanum* | 1 | 0 | 0 | 0 |
| *Sorghum bicolor* | 7 | 1 | 1 | 0 |
| *Secale cereal* | 1 | 0 | 0 | 0 |
| *Setaria italica* | 7 | 1 | 1 | 0 |
| *Spirodela polyrhiza* | 5 | 0 | 1 | 1 |
| *Triticum aestivum* | 7 | 1 | 0 | 0 |
| *Triticum monococcum* | 1 | 0 | 0 | 0 |
| *Zantedeschia aethiopica* | 3 | 0 | 0 | 0 |
| *Zostera marina* | 4 | 1 | 1 | 1 |
| *Zea mays* | 9 | 1 | 1 | 0 |
| **Eudicots** |  |  |  |  |
| *Aquilegia coerulea* | 2 | 1 | 0 | 0 |
| *Aquilegia formosa x Aquilegia pubescens* | 4 | 1 | 0 | 0 |
| *Arabidopsis halleri* | 0 | 1 | 0 | 0 |
| *Arabidopsis lyrata* | 8 | 1 | 1 | 0 |
| *Arabidopsis thaliana* | 6 | 1 | 1 | 0 |
| *Arachis hypogaea* | 2 | 0 | 0 | 0 |
| *Boechera stricta* | 2 | 0 | 0 | 0 |
| *Brassica juncea* | 2 | 0 | 0 | 0 |
| *Brassica napus* | 4 | 1 | 0 | 0 |
| *Brassica oleracea* | 2 | 1 | 0 | 0 |
| *Brassica rapa* | 2 | 1 | 0 | 0 |
| *Camellia sinensis* | 1 | 0 | 0 | 0 |
| *Capsella grandiflora* | 6 | 1 | 1 | 0 |
| *Capsicum annuum* | 4 | 0 | 0 | 0 |
| *Carica papaya* | 3 | 0 | 0 | 0 |
| *Catharanthus roseus* | 4 | 1 | 0 | 0 |
| *Cichorium intybus* | 1 | 0 | 0 | 0 |
| *Citrus clementina* | 5 | 1 | 1 | 0 |
| *Citrus sinensis* | 5 | 1 | 0 | 0 |
| *Codonopsis lanceolata* | 1 | 0 | 0 | 0 |
| *Cucumis melo* | 1 | 0 | 0 | 0 |
| *Cucumis sativus* | 5 | 1 | 1 | 0 |
| *Cucurbita* cv. Kurokawa Amakuri | 3 | 0 | 0 | 0 |
| *Eucalyptus camaldulensis* | 3 | 1 | 0 | 0 |
| *Eucalyptus globulus* | 6 | 1 | 1 | 0 |
| *Eucalyptus grandis* | 6 | 1 | 1 | 0 |
| *Eucalyptus gunnii* | 6 | 1 | 1 | 0 |
| *Fragaria x ananassa* | 3 | 0 | 0 | 0 |
| *Gerbera hybrida* cv. Terra Regina | 2 | 0 | 0 | 0 |
| *Glycine max* | 7 | 1 | 2 | 1 |
| *Gossypium hirsutum* | 2 | 1 | 0 | 0 |
| *Gossypium raimondii* | 3 | 1 | 0 | 0 |
| *Hedyotis centranthoides* | 1 | 0 | 0 | 0 |
| *Hedyotis terminalis* | 1 | 0 | 0 | 0 |
| *Helianthus annuus* | 5 | 1 | 0 | 0 |
| *Helianthus argophyllus* | 2 | 0 | 0 | 0 |
| *Helianthus paradoxus* | 1 | 0 | 0 | 0 |
| *Hevea brasiliensis* | 2 | 0 | 0 | 0 |
| *Ipomoea batatas* | 1 | 0 | 0 | 0 |
| *Ipomoea nil* | 2 | 0 | 0 | 0 |
| *Lactuca sativa* | 2 | 0 | 0 | 1 |
| *Linum usitatissimum* | 5 | 1 | 2 | 1 |
| *Lotus japonicus* (corniculatus var japonicus) | 2 | 0 | 0 | 0 |
| *Lycopersicon esculentum* | 8 | 1 | 1 | 1 |
| *Malus domestica* | 2 | 1 | 0 | 0 |
| *Manihot esculenta* | 7 | 1 | 1 | 0 |
| *Medicago truncatula* | 5 | 1 | 1 | 0 |
| *Mesembryanthemum crystallinum* | 4 | 2 | 0 | 1 |
| *Mimulus guttatus* | 4 | 1 | 1 | 0 |
| *Nicotiana tabacum* | 6 | 0 | 2 | 1 |
| *Pimpinella brachycarpa* | 1 | 0 | 0 | 0 |
| *Pisum sativum* | 1 | 0 | 0 | 0 |
| *Poncirus trifoliata* | 2 | 0 | 0 | 0 |
| *Populus balsamifera* | 3 | 0 | 0 | 0 |
| *Populus tomentosa* | 1 | 0 | 0 | 0 |
| *Populus trichocarpa* | 8 | 1 | 1 | 0 |
| *Prunus persica* | 5 | 1 | 1 | 0 |
| *Quercus shumardii* | 1 | 0 | 0 | 0 |
| *Raphanus sativus* | 1 | 0 | 0 | 0 |
| *Rehmannia glutinosa* | 1 | 0 | 0 | 0 |
| *Retama raetam* | 1 | 0 | 0 | 0 |
| *Rheum australe* | 1 | 0 | 0 | 0 |
| *Ricinus communis* | 2 | 1 | 0 | 2 |
| *Solanum tuberosum* | 5 | 0 | 0 | 0 |
| *Spinacia oleracea* | 3 | 0 | 0 | 1 |
| *Suaeda salsa* | 1 | 0 | 0 | 0 |
| *Tarenaya hassleriana* | 0 | 1 | 0 | 0 |
| *Thellungiella parvula* | 1 | 0 | 1 | 0 |
| *Thellungiella salsuginea* | 9 | 1 | 0 | 0 |
| *Vigna unguiculata* | 4 | 0 | 0 | 0 |
| *Vitis hybrid cultivar* | 1 | 0 | 0 | 0 |
| *Vitis vinifera* | 4 | 2 | 1 | 0 |

**Table S2.**
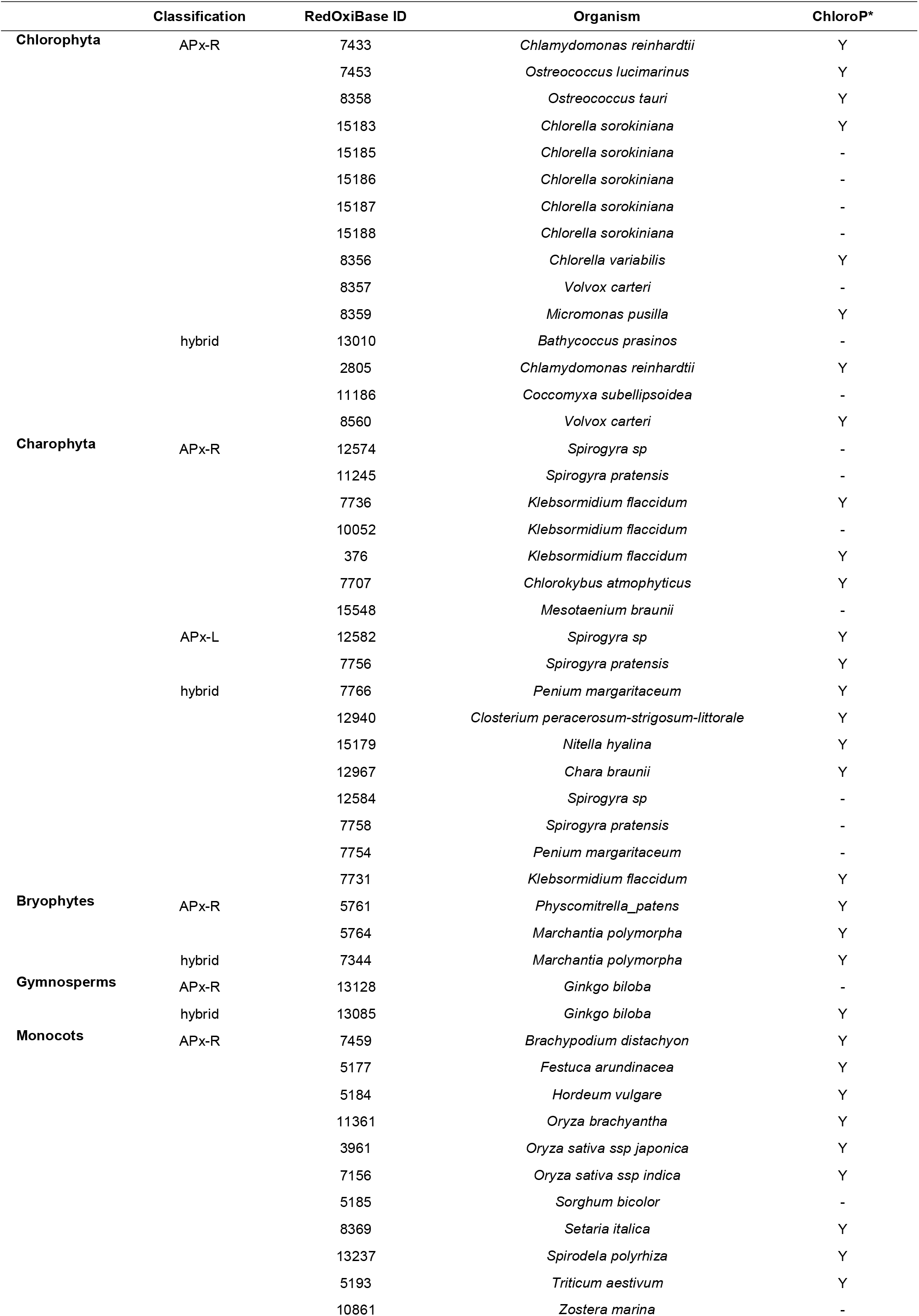
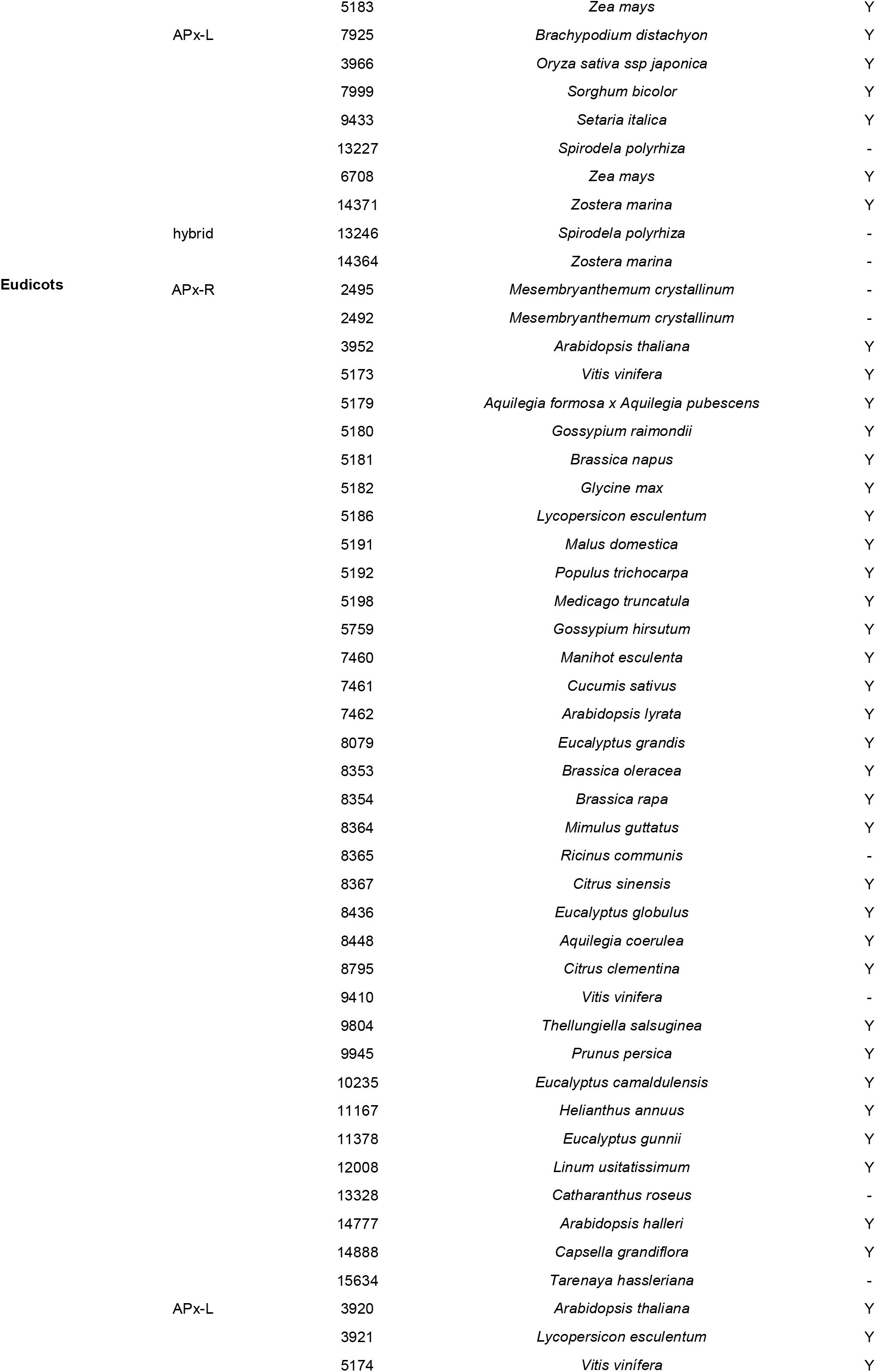
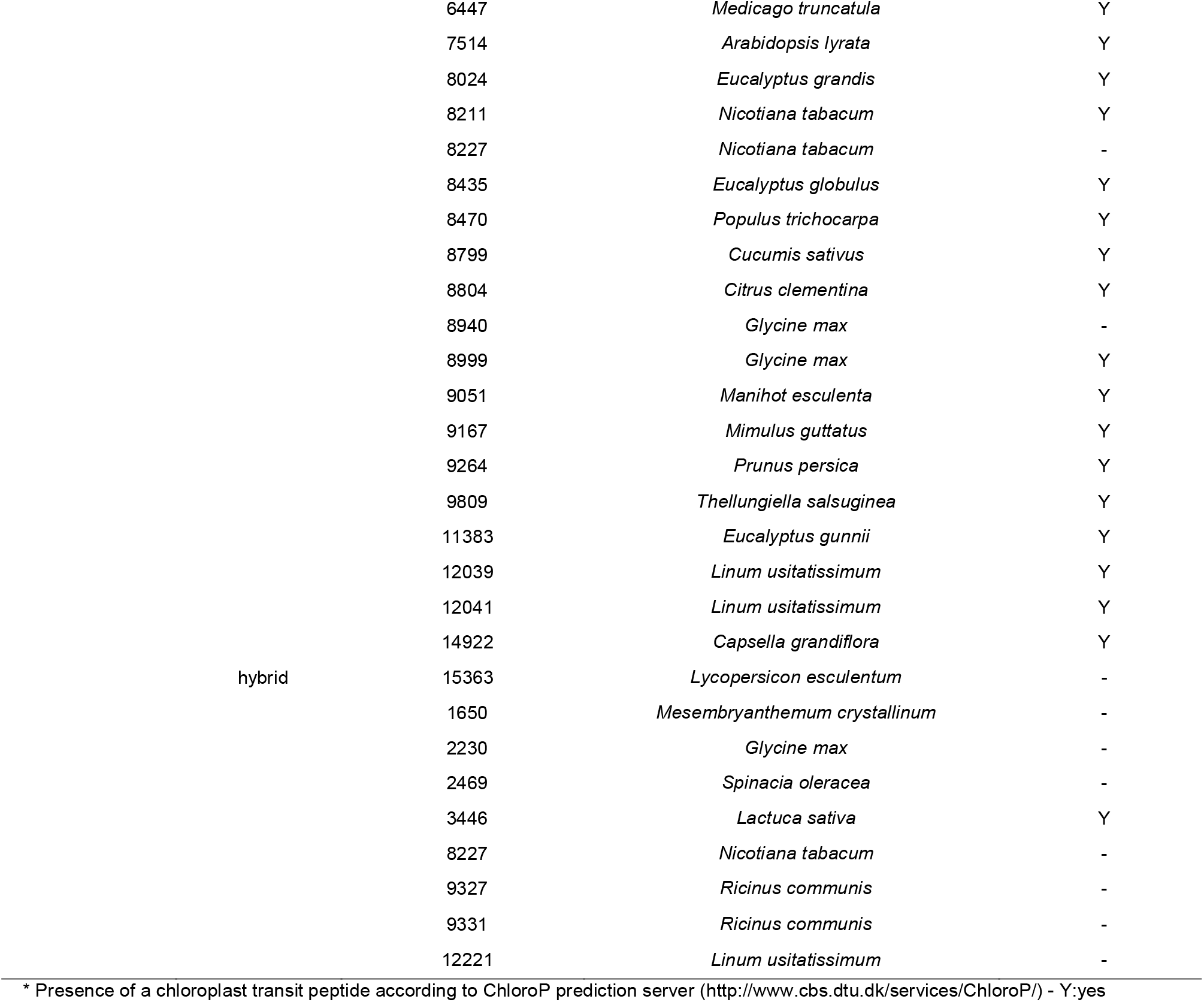
Accession numbers of APx-R, APx-L and hybrid proteins deposited in RedOxiBase.

**Table S3**. Charophytes sequences retrieved from OneKP used in the analyses.

## Acknowledgements

This work was supported by Coordenação de Aperfeiçoamento de Pessoal de Nível Superior (CAPES) and Conselho Nacional de Desenvolvimento Científico e Tecnológico (CPNq).

## References

1 Shigeoka S, Ishikawa T, Tamoi M, Miyagawa Y, Takeda T, Yabuta Y & Yoshimura K (2002) Regulation and function of ascorbate peroxidase isoenzymes. J. Exp. Bot. 53, 1305–1319.

2 Lad L, Mewies M & Raven EL (2002) Substrate binding and catalytic mechanism in ascorbate peroxidase: Evidence for two ascorbate binding sites. Biochemistry 41, 13774–13781.

3 Patterson WR & Poulos TL (1995) Crystal Structure of Recombinant Pea Cytosolic Ascorbate Peroxidase. Biochemistry 34, 4331–4341.

4 Mandelman D, Li H, Poulos TL & Schwarz FP (2009) The role of quaternary interactions on the stability and activity of ascorbate peroxidase. Protein Sci. 7, 2089–2098.

5 Raven EL (2003) Understanding functional diversity and substrate specificity in haem peroxidases: What can we learn from ascorbate peroxidase? Nat. Prod. Rep. 20, 367–381.

6 Sharp KH, Mewies M, Moody PCE & Raven EL (2003) Crystal structure of the ascorbate peroxidase-ascorbate complex. Nat. Struct. Biol. 10, 303–307.

7 Macdonald IK, Badyal SK, Ghamsari L, Moody PCE & Raven EL (2006) Interaction of ascorbate peroxidase with substrates: A mechanistic and structural analysis. Biochemistry 45, 7808–7817.

8 Kovacs FA, Sarath G, Woodworth K, Twigg P & Tobias CM (2013) Abolishing activity against ascorbate in a cytosolic ascorbate peroxidase from switchgrass. Phytochemistry 94, 45–52.

9 Çelik A, Cullis PM, Sutcliffe MJ, Sangar R & Raven EL (2001) Engineering the active site of ascorbate peroxidase. Eur. J. Biochem. 268, 78–85.

10 Battistuzzi G, Bellei M, Bortolotti CA & Sola M (2010) Redox properties of heme peroxidases. Arch. Biochem. Biophys. 500, 21–36.

11 Pipirou Z, Bottrill AR, Metcalfe CM, Mistry SC, Badyal SK, Rawlings BJ & Raven EL (2007) Autocatalytic formation of a covalent link between tryptophan 41 and the heme in ascorbate peroxidase. Biochemistry 46, 2174–2180.

12 Banci L (1997) Structural properties of peroxidases. J. Biotechnol. 53, 253–263.

13 Chary KVR & Srivastava AK (2013) Encyclopedia of biophysics.

14 Welinder K G (1992) Superfamily of plant, fungal and bacterial peroxidases. Curr. Opin. Struct. Biol. 2, 388–393.

15 Passardi F, Bakalovic N, Teixeira FK, Margis-Pinheiro M, Penel C & Dunand C (2007) Prokaryotic origins of the non-animal peroxidase superfamily and organelle-mediated transmission to eukaryotes. Genomics 89, 567–579.

16 Zámocký M, Furtmüller PG & Obinger C (2010) Evolution of structure and function of Class I peroxidases. Arch. Biochem. Biophys. 500, 45–57.

17 Lazzarotto F, Turchetto-Zolet AC & Margis-Pinheiro M (2015) Revisiting the Non-Animal Peroxidase Superfamily. Trends Plant Sci. 20, 807–813.

18 Lazzarotto F, Teixeira FK, Rosa SB, Dunand C, Fernandes CL, de Vasconcelos Fontenele A, Silveira JAG, Verli H, Margis R & Margis-Pinheiro M (2011) Ascorbate peroxidase-related (APx-R) is a new heme-containing protein functionally associated with ascorbate peroxidase but evolutionarily divergent. New Phytol. 191, 234–250.

19 Chen C, Letnik I, Hacham Y, Dobrev P, Ben-Daniel B-H, Vanková R, Amir R & Miller G (2014) ASCORBATE PEROXIDASE6 Protects Arabidopsis Desiccating and Germinating Seeds from Stress and Mediates Cross Talk between Reactive Oxygen Species, Abscisic Acid, and Auxin. Plant Physiol. 166, 370–383.

20 Lundberg E, Storm P, Schröder WP & Funk C (2011) Crystal structure of the TL29 protein from Arabidopsis thaliana: An APX homolog without peroxidase activity. J. Struct. Biol. 176, 24–31.

21 Granlund I, Storm P, Funk C, Schröder WP, García-Cerdán JG, Schubert M & Granlund I (2009) The TL29 Protein is Lumen Located, Associated with PSII and Not an Ascorbate Peroxidase. Plant Cell Physiol. 50, 1898–1910.

22 Wang Y-Y, Hecker AG & Hauser BA (2014) The APX4 locus regulates seed vigor and seedling growth in Arabidopsis thaliana. Planta 239, 909–919.

23 Lazzarotto F, Turchetto-Zolet AC & Margis-Pinheiro M (2015) Revisiting the Non-Animal Peroxidase Superfamily. Trends Plant Sci. 20.

24 Poulos TL (2011) Thirty Years of Heme Peroxidase Structural Biology. Arch Biochem Biophys. 500, 3–12.

25 Lad L, Mewies M, Basran J, Scrutton NS & Raven EL (2002) Role of histidine 42 in ascorbate peroxidase: Kinetic analysis of the H42A and H42E variants. Eur. J. Biochem. 269, 3182–3192.

26 Kitajima S, Kitamura M & Koja N (2008) Triple mutation of Cys26, Trp35, and Cys126 in stromal ascorbate peroxidase confers H2O2 tolerance comparable to that of the cytosolic isoform. Biochem. Biophys. Res. Commun. 372, 918–923.

27 Yergalem T. Meharenna, Patricia Oertel, B. Bhaskar and TLP (2008) Engineering Ascorbate Peroxidase Activity into Cytochrome c Peroxidase. Biochemistry 47, 10324–10332.

28 Teixeira FK, Menezes-Benavente L, Margis R & Margis-Pinheiro M (2004) Analysis of the molecular evolutionary history of the ascorbate peroxidase gene family: Inferences from the rice genome. J. Mol. Evol. 59, 761–770.

29 Turner DD, Lad L, Kwon H, Basran J, Carr KH, Moody PCE & Raven EL (2018) The role of Ala134 in controlling substrate binding and reactivity in ascorbate peroxidase. J. Inorg. Biochem. 180, 230–234.

30 Savelli B, Li Q, Webber M, Jemmat AM, Robitaille A, Zámocký M, Mathé C & Dunand C (2019) RedoxiBase: A database for ROS homeostasis regulated proteins. Redox Biol. 26, 0–4.

31 Edgar RC, Drive RM & Valley M (2004) MUSCLE_: multiple sequence alignment with high accuracy and high throughput. 32, 1792–1797.

32 Kumar S, Stecher G & Tamura K (2016) MEGA7_: Molecular Evolutionary Genetics Analysis Version 7. 0 for Bigger Datasets Brief communication. 33, 1870–1874.

33 Vaughan T, Wu C, Xie D, Suchard MA, Rambaut A & Drummond AJ (2014) BEAST 2_: A Software Platform for Bayesian Evolutionary Analysis. 10, 1–6.

34 Abascal F, Zardoya R & Posada D (2005) ProtTest_: selection of best-fit models of protein evolution. 21, 2104–2105.

35 Gilks WR (2005) Markov Chain Monte Carlo. Encycl. Biostat., 1–7.

36 Rambaut A, Drummond AJ, Xie D, Baele G & Suchard MA (2018) Posterior Summarization in Bayesian Phylogenetics Using Tracer 1. 7. 67, 901–904.

37 Hu B, Jin J, Guo AY, Zhang H, Luo J & Gao G (2015) GSDS 2.0: An upgraded gene feature visualization server. Bioinformatics 31, 1296–1297.

